# Phage resistance impacts antibiotic susceptibility and virulence in *Staphylococcus aureus*

**DOI:** 10.1101/2025.09.02.673687

**Authors:** Janine Bowring, Freja C. Mikkelsen, Roshni Haider, Esther Lehmann, Thibault Frisch, Morten Kjos, Nina M. van Sorge, Hanne Ingmer

## Abstract

As antibiotic resistance continues to rise worldwide, phage therapy offers a promising solution by harnessing viruses that infect and kill bacteria. However, phage-resistance may develop, compromising therapy. Phage-resistance has primarily been linked to changes in major phage receptors that for the human pathogen *Staphylococcus aureus* are the cell-wall linked wall teichoic acids (WTA). To identify additional factors contributing to phage-resistance, *S. aureus* was evolved under selective pressure from lytic phages, including candidates from therapeutic phage cocktails, to obtain resistant clones. A third of the phage-resistant clones acquired mutations associated with cell-wall changes previously linked with phage resistance, namely in *femA*, involved in peptidoglycan crossbridge formation, and *tagO*, encoding the initiator of the WTA biosynthesis. The remaining clones had mutations in pathways not previously associated with phage-resistance, including nucleoside catabolism (*deoC1*), polyamine import (*potAB*) and DNA/RNA replication (*cshA, ligA*), suggesting that phage-resistance may be associated with phage-driven host takeover. Antibiotic susceptibility and virulence were diversely affected by the mutations; mutations in *tagO* increased β-lactams sensitivity and attenuated virulence, whereas mutations in *femA*/*ligA* or *cshA* increased staphylococcal virulence in *Galleria mellonella*. This work demonstrates the involvement of several novel bacterial pathways in phage resistance, highlighting the complexity behind phage-bacterial interactions.

## Introduction

The combination of antibiotics and bacterial viruses, known as bacteriophages or phages, is of growing interest for treatment of infections with antibiotic-resistant pathogens. One such pathogen is *Staphylococcus aureus* that can cause a variety of infections (1). *S. aureus* is known for its resistance to cell wall acting antibiotics, including β-lactams in methicillin resistant *S. aureus* (MRSA) strains and vancomycin in the vancomycin intermediate resistant (VISA) strains. For treatment of staphylococcal infections, several phages are commercially available and have been used with success in a growing number of patients (2). However, currently we have a limited understanding of the factors contributing to phage resistance and how phage resistance impacts bacterial properties, such as virulence and antibiotic susceptibility.

The primary phage receptor in *S. aureus* is the wall teichoic acid (WTA), a glycopolymer covalently linked to the peptidoglycan cell wall (3, 4). WTA synthesis is initiated by TagO (5) after which a polymer consisting of ribitol phosphates is synthesized (6) that is further modified by N-acetylglucosamine (GlcNAc). WTA GlcNAc attachment is catalyzed by the glycosyltransferases TarM (α-1,4-GlcNAc) and TarS (β-1,4-GlcNAc), as well as the prophage encoded TarP (β-1,3-GlcNAc). Interestingly, the glycosylation pattern of WTA impacts phage susceptibility; K-type myoviruses bind to the WTA backbone whereas others require either ⍺- and/or β-GlcNAc modifications for binding and subsequent infection (4). Recently, we showed that the *S. aureus agr* quorum sensing system controls the WTA α/β-GlcNAc glycosylation pattern and phage susceptibility by suppressing *tarM* expression in stationary phase (7). Also, the ArlRS two-component system has been shown to modulate the WTA α/β-GlcNAc glycosylation pattern affecting phage susceptibility (8). However, less is known about intracellular factors protecting *S. aureus* against phages, with knowledge mainly limited to the presence or absence of phage defense systems, including restriction modification systems, phage inducible chromosomal islands, and CRISPR-Cas (9). However, some defense systems offer limited protection against lytic phages in *S. aureus* (10), and susceptibility to lytic phages has been primarily linked to altered phage receptors (11, 12).

Intriguingly, WTA is also linked to susceptibility to cell wall targeting antibiotics. In MRSA, WTA β-GlcNAc but not α-GlcNAc was shown to be involved in β-lactam (13), methicillin (14), daptomycin (15) and vancomycin (16) resistance. Therefore, loss of WTA or alterations of WTA glycosylation patterns may affect both phage and antibiotic susceptibility.

Several studies have identified synergism between *S. aureus* lytic phages and β-lactam antibiotics, including cefoxitin, oxacillin, and ampicillin (17-19). One recent evolution experiment, where *S. aureus* was exposed to the K-type lytic virus ϕStaph1N, identified phage resistant clones with increased susceptibility to β-lactams (20). A similar experiment using the lytic phage PYO_sa_ resulted in reduced phage susceptibility of small colony variants with mutations in *femA*, encoding a protein catalyzing the formation of pentaglycine cross-bridges in the peptidoglycan layer of the cell wall (21). In this case, synergy with antibiotics only occurred with sequential treatment of phages and then antibiotics, while simultaneous treatment reduced the effectiveness of PYO_sa_ killing.

It has been shown that WTA is important for *S. aureus* immune interactions (22), endothelium adhesion (23) and nasal colonization (24) through interactions with human immune receptors, scavenger receptors, and soluble receptors in serum (9). WTA has been shown to activate CD4+ T cells through the major histocompatibility complex II, inducing abscess formation (25), while WTA is also required for release of cytolytic toxins (26). Indeed, highly pathogenic strains of *S. aureus* often have elevated WTA content in their cell wall compared to less virulent strains, and overexpression of WTA can be a mechanism for *S. aureus* to gain virulence (27).

Here, we generated *S. aureus* mutants resistant to three well-known K-type myoviruses, namely phage K (28), ϕIPLA-RODI (29) and Stab21 (30), that all have been proposed as candidates for phage-based therapeutics. These phages are characterized by their myovirus morphology and lytic activity against *S. aureus*, with >80% genome identity between their genomes and all three binding to the WTA backbone. K-type myoviruses carry 2 receptor binding proteins (RBPs), where RBP1 preferentially binds glycosylated WTA and RBP2 preferentially binds unglycosylated RboP-WTA (31). We characterized evolved phage-resistant mutants genotypically and confirmed their resistance phenotypes by CRISPRi knock-down of the identified genes. Phenotypic characterizations included antibiotic susceptibility testing to β-lactams and vancomycin, WTA glycosylation profiles, and virulence potential in an *in vivo Galleria mellonella* infection model. Our results show that decreased phage susceptibility is associated with mutations in multiple genes, including mutations in genes not previously linked to phage resistance. In contrast to previous findings where phage resistance resulted in increased β-lactam susceptibility (20), we show that phage resistance can be associated with both increased and decreased β-lactam susceptibility, highlighting the complexity underlying *S. aureus* susceptibility to phages and antibiotics.

## Materials and methods

### Bacterial strains and phages

Bacterial strains used in this study are shown in Table S1. *S. aureus* strains were cultured in Tryptone Soy Broth (TSB, Oxoid) or on Tryptone Soy Agar (TSA, Oxoid) at 37°C. For CRISPRi strains, media was supplemented with antibiotics spectinomycin (spec, 250 µg mL^-1^, Fisher Scientific, 15480207) and chloramphenicol (cam, 10 µg mL^-1^, Sigma, C0378), as well as IPTG (250 µg mL^-1^, Fisher Scientific, R0392) where appropriate. Phages are detailed in Table S2 and were propagated as described in (32).

### Generation of bacteriophage resistant mutant library

Wild-type *S. aureus* USA300 JE2 was exposed to one of three well-characterized lytic staphylococcal phages (Fig 1A; Table S2). An overnight culture of JE2 was diluted to OD_600_ 0.05 and grown at 37°C with shaking to OD_600_ 0.35 (≅5 x 10^7^ CFU mL^-1^). 100 µl of the culture was mixed with 100 µl of phage at a multiplicity of infection (MOI) of 1. Samples were incubated at room temperature for 10 min before 3 mL molten phage top agar (PTA; Oxoid nutrient broth no. 2, agar 3.5% wt/vol) with 10mM CaCl_2_ was added. The mixture was plated on phage base agar plates (PBA: Oxoid nutrient broth no. 7, agar 3.5% wt/vol) with 10mM CaCl_2_ and incubated overnight at 37°C. Most of the bacteria were lysed and growth appeared as individual colonies, which were re-streaked twice on TSA plates before storage at -80°C. This step was performed to select for resistance through mutation rather than expression changes or small colony variants, and to remove phages potentially carried over from the surrounding plate.

**Figure 1.**
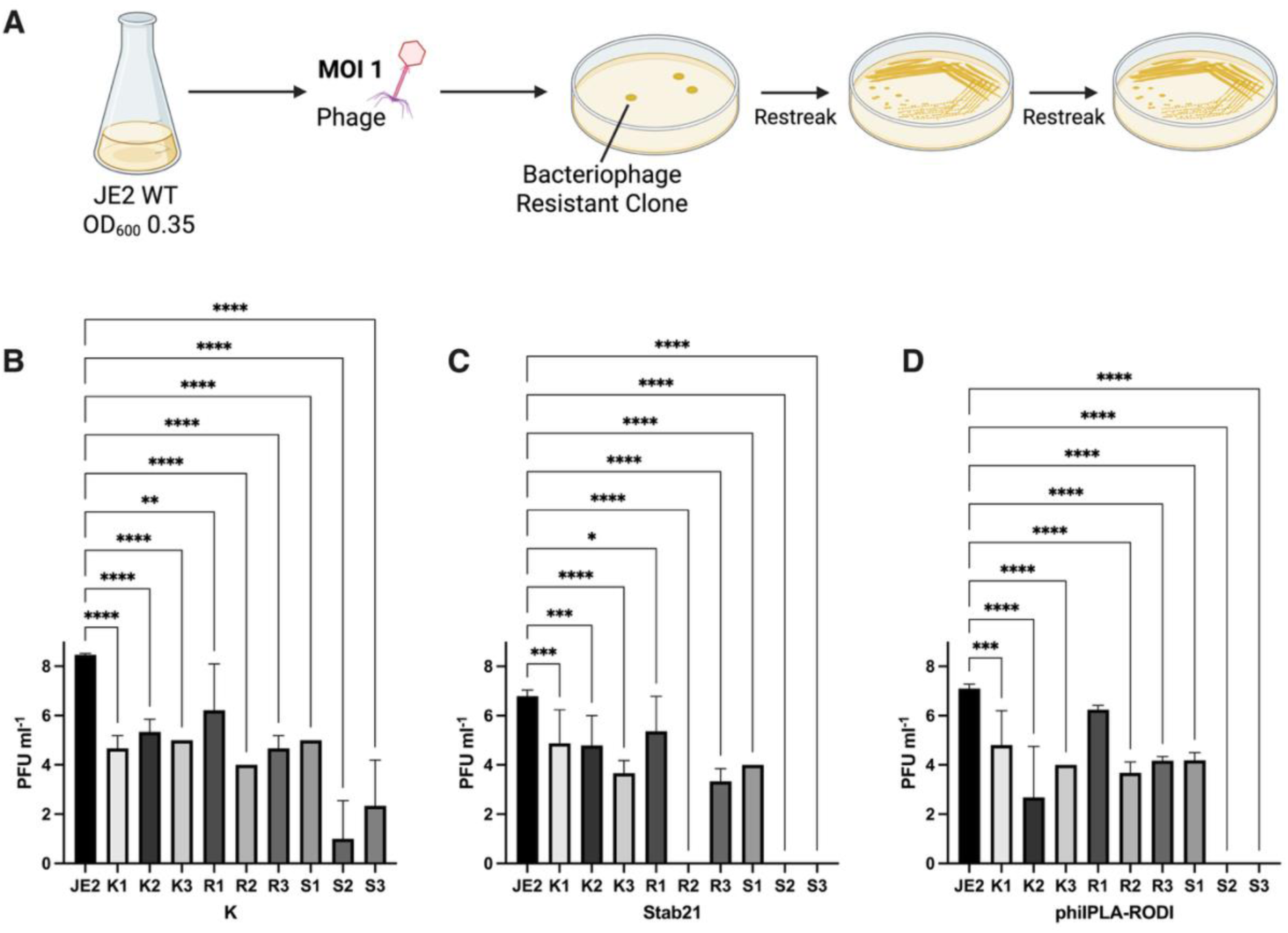
Evolving bacteriophage resistant *S. aureus* clones. **A.** Workflow of the bacteriophage resistant clone development protocol. **B.** PFU ml^-1^ of phage K C. Stab21 D. ϕIPLA-RODI on the phage resistant clones compared to the parental JE2 strain. For B-D, data shows mean and standard deviation of 6 biological replicates that was log transformed. Statistical analysis was performed using ordinary one-way ANOVA, with multiple comparisons to compare each mutant to the JE2, with Dunnett’s multiple comparisons test. Significant differences are indicated by one (*p* <0.05), two (*p* <0.01), three (*p* <0.001) or four (*p* <0.0001) asterisks (*).

### Phage titer assay

Plaque forming units per mL (PFU mL^-1^) were assessed as previously described (32). Briefly, *S. aureus* overnight cultures were diluted to OD_600_ 0.05 and grown at 37°C with shaking to OD_600_ 0.35. 100 µL of culture was mixed in 3 mL molten PTA with 10 mM CaCl_2_ and plated on PBA with 10 mM CaCl_2_. Phage lysates were diluted 10-fold in phage buffer (1 mM MgSO_4_, 4 mM CaCl_2_, 50 mM Tris-HCl [pH: 8], 0.1 M NaCl) and spotted on the bacterial lawn (10 µL per spot; dilutions 10^0^-10^-7^). Plaques were enumerated after overnight incubation at 37°C.

### Antibiotic susceptibility assay

The minimum inhibitory concentrations (MIC) of different *S. aureus* strains for a range of antibiotics were measured on Mueller-Hinton agar plates (Oxoid), using Etests (BioMérieux) according to the manufacturer’s guidelines. 2% NaCl was added to the agar for oxacillin Etests. Briefly, freshly streaked colonies were resuspended in saline to a McFarland standard of 0.5 and streaked on the MH agar plate. Plates were dried before adding the Etest strips and incubating overnight at 37°C. Four cell wall targeting antibiotics were tested: vancomycin (VAN, 256 µg ml^-1^), cefoxitin (FOX, 256 µg ml^-1^), cefotaxime (CTX, 32 µg ml^-1^) and oxacillin (OXA, 256 µg ml^-1^). The MIC was read at the concentration where the bacterial growth intersected the Etest.

### Whole genome sequencing and mutation analysis

DNA of the parental strain USA300 JE2 and the phage-resistant mutants was extracted using the GenElute™ Bacterial Genomic DNA Kit (Merck), as per manufacturer’s instructions for Gram-positive bacteria. Whole genome sequencing (WGS) was performed by Illumina MiSeq (Eurofins). Sequence analysis was performed in Geneious Prime (v2025.1) where sequences were trimmed for adapters and quality with the BBDuk plugin and aligned to the NCBI USA300 JE2 reference sequence (CP020619.1) using the Geneious mapper. Variations and single nucleotide polymorphisms (SNPs) were identified using the Geneious ‘Find Variations/SNPs’ tool and phage resistant strain SNPs were compared to the parental JE2 sequencing. A cut off of 50% frequency and 30 read coverage was used for the final SNP Table (Table 1).

**Table 1.**
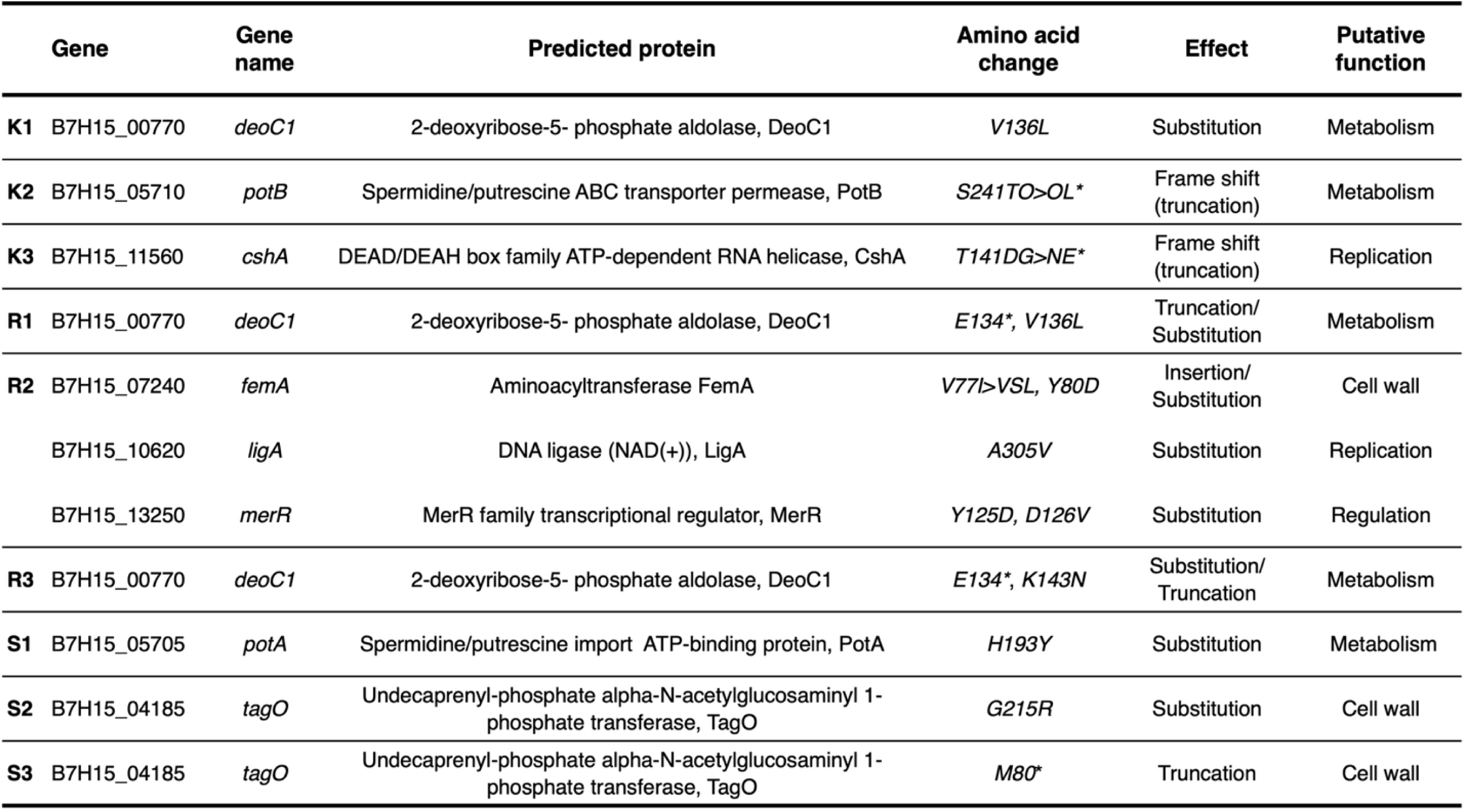
SNPs identified by whole genome sequencing in the phage resistant *S. aureus* clones. Putative functions were categorized by COG and PFAM NCBI databases. Naming system indicates which phage the clone was exposed to during evolution (K, phage K; R, ϕIPLA-RODI; S, Stab21). *\** indicates stop codon.

### Correlation analysis

The relationship between phage and antibiotic susceptibility was assessed using a correlation matrix with Pearson correlation coefficients in GraphPad Prism (v. 10.4). The PFU ml^-1^ was normalized by log transformation. To make the log(PFU) and MIC results comparable in continuous format, the log(PFU) was inverted by 1/log(PFU) so that for both data sets higher values indicated higher resistance. The Z-scores for all strains were calculated using SciPy (v1.15.2) (33) to ensure comparability between datasets and plotted in the correlation matrix.

### Hemolysis phenotype and quantification

Hemolysis phenotyping was performed by streaking, dotting, or cross-streaking *S. aureus* on TSA blood plates (TSA, Oxoid; 5% sheep blood, Hatunalab). For dot screening, target strains were transferred from a streaked plate in dots to investigate α, β or γ hemolysis (34), while to investigate for δ hemolysis cross streaking was conducted as described in (35), using *S. aureus* RN4220 for the central streak. For quantification of overall hemolysis, target strains were grown from a single colony overnight and diluted to OD_600_ 0.02 before growing overnight again, whereafter 1 mL of each sample was spun down at 6000 rpm for 1 min and 200 µL of the supernatant was collected. The supernatants were mixed with 775 µL PBS and 25 µL defibrinated sheep blood (Hatunalab AB) and incubated at 37°C for 1h. Samples were then spun down at 3000 rpm for 10 min and OD_450_ was measured to indicate hemolysis effectivity and compared to positive (200µl 1% Triton X-100) and negative (200µl TSB) controls.

### WTA anti-GlcNAc-antibody deposition assay

Table S3 contains a list of all antibodies used in this study. Antibody staining for *S. aureus* WTA was performed similarly to (36), using anti-α1,4-GlcNAc-WTA Fab (clone 4461) and anti-β-,4-GlcNAc-WTA Fab (clone 4462) (37) to avoid epitope-independent interaction with staphylococcal protein A. In brief, bacterial cultures were adjusted to 1.25 x 10^6^ cells in PBS-0.1% BSA (1x phosphate-buffered saline with 0.1% bovine serum albumin; Sigma A7030) and incubated with the Fab-fragments (3-fold serial dilution from 10 µg/mL - 0.1 µg/mL) for 30 min at 4°C. After washing with PBS-0.1% BSA, bacteria were incubated with 1:1000 diluted Goat F(ab’)2 anti-human-kappa-AF647 secondary antibody (SouthernBiotech, Cat. No.: 2062-31) in PBS-0.1% BSA for 20 min at 4°C. Unbound antibodies were removed, and bacteria were fixed in 1% formaldehyde in 1x PBS for 15 min. Flow cytometry data were acquired on an FACSCanto (BD) using FACSDiva (BD). Per sample, 10,000 events were measured within the set gate. All data were analyzed using FlowJo 10 (FlowJo, LLC).

### Growth curves

Growth curves were performed using a plate reader (Bioscreen C, Type FP-1100-C) at 37°C and 200 rpm for 24 h. Overnight cultures were adjusted to 0.05 OD_600_ and OD_600_ was measured at 20 min intervals. Shaking was halted 5 s before measurement. Growth curves were analyzed using GrowthCurver (v0.3.1) (38) in RStudio (2024.12.0+467; R version 4.4.2) to fit a logistic curve and calculate the generation time (t_gen) and area under the curve (AUC).

### Galleria mellonella infection model

Virulence of the JE2 wild type (wt) and selected phage-resistant mutants (R2, S2, S3, K3, Δ*tagO*) was evaluated in a *Galleria mellonella* intrahemocoelic injection model. Larvae were acquired through a local pet shop (www.krybdyrsiden.dk) and, if not used directly, kept at 15°C. The day before infection, larvae were separated into groups of ten and kept overnight at 37°C. Larvae were infected with 10^6^ CFU (bacteria washed thrice in 1x PBS and CFU adjusted with 1x PBS) in a total volume of 10 µL through the front right pro-leg using an automatic injector setup (Hamilton PB600-1 Repeating Syringe Dispenser combined with Hamilton 250 µL, Gastight Syringe (RN) and needle type AS). To monitor basic larvae health, we included an uninfected injection control group (1x PBS), as well as an uninjected control group. Following infection larvae were kept at 37°C and their survival was monitored for 60 h. Health was scored in 12 h intervals using “alive” and “dead” as scoring criteria.

### CRISPRi knockdown of potential phage resistance genes

Knockdown constructs were designed through single guide RNAs targeting the 20 bp sequence adjacent to the 5’ proximal PAM on the non-template strain for genes of interest. These were selected and cloned into the pVL2336 plasmid, as previously described (39). Strain JE2 was transformed with both the sgRNA plasmids and an IPTG-inducible pLOW plasmid encoding the dead cas9 (pLOW-*dcas9* aad9). For the knockdown phage resistance assay, strains were grown overnight from a single colony, before diluting to OD_600_ 0.05. They were then grown for 2.5 h at 37°C with 180 rpm, before subculturing 1/50 in 125 μL TSB plus 500 μM IPTG for 1 h at 37°C with 180 rpm shaking. After 1 h, 125 μL of phage K (MOI 10^-4^ or 10^-5^) or phage buffer was added (250 μM final IPTG). OD_600_ was recorded every 20 min for 16 h at 30°C with shaking using a plate reader (Bioscreen C, Type FP-1100-C). Phage resistance was defined as an IPTG induced delay in phage killing compared to the non-induced control.

## Results

### Generation of bacteriophage resistant clones

A library of bacteriophage-resistant clones was generated in the MRSA strain USA300 JE2 (JE2), using an adaptation of the classic bacteriophage overlay assay (Figure 1A) with one of three lytic phages: phage K, ϕIPLA-RODI, or Stab21. Surviving colonies were assessed for phage susceptibility compared to JE2 (Figure S1, n = 14). Of these, nine phage resistant clones were selected for further studies and labelled according to the phage against which resistance was developed with “K” signifying phage K, “R” for ϕIPLA-RODI and “S” for Stab21. Compared to the JE2 parent strain, these strains displayed statistically significant reductions in susceptibility to the phage against which they had been evolved (PFU ml^-1^). Other phenotypes including growth, colony morphology and hemolysis remained similar to JE2 (Fig S2A-F). Two exceptions were S2 and S3, which showed attenuated growth, smaller colonies and lower levels of hemolysis.

The nine selected clones were characterized further for susceptibility to all three phages (Fig 1B-D). Clones S2 and S3 were completely resistant to Stab21 (Fig 1D) and ϕIPLA-RODI (Fig 1C) infections, while occasionally unclear plaques formed with phage K when exposing the bacteria to phage concentrations of 10^7^ to 10^8^ PFU ml^-1^ (Fig 1B). Clone R2 had a 3-log reduction in susceptibility to ϕIPLA-RODI for which it was selected and displayed a 4-log reduction for phage K but was completely resistant to Stab21. Clone R1 displayed only a minor decrease in susceptibility to the three different phages (1 to 2 log reduction in PFU ml^-1^), while the other clones (K1-K3, R3, and S1) showed intermediate levels of phage resistance (between 3 to 5 log reduction in PFU ml^-1^ towards all phages).

### Phage-resistant clones acquired mutations in cell wall synthesis, metabolism, replication, and regulation-associated genes

To determine the genotypes behind the changes in phage susceptibility, whole genome sequencing with variation/SNP analyses was performed. Mutations were located in genes associated with cell wall synthesis, transcriptional regulation, transport and metabolism, or replication recombination and repair, when classified using Clusters of Orthologous Genes (COG) (40) and Conserved Protein Domain Family (PFAM) (41) NCBI databases (Table 1; Table S4). Eight out of nine clones were mutated in a single gene, whereas one clone (R2) had mutations in three genes (Table 1). The two clones that displayed the most profound phage resistance, S2 and S3, had a SNP or truncating mutation in *tagO,* respectively (Table 1), supporting previous findings that *tagO* is essential for *S. aureus* phage infection (42). The S2 *tagO* SNP (G215R) is located centrally in an AlphaFold predicted protein structure (Fig S3) and changes the side chain size, charge and polarity. Therefore, it is likely this SNP and the truncation in S3 (M80*) would render the TagO enzyme inactive and disrupt WTA synthesis. The cell wall was also affected in clone R2, which had substitutions in *femA, ligA,* and *merR*. LigA is a DNA ligase involved in replication (43) and *merR* encodes a transcriptional regulator (44), while FemA catalyzes the formation of the pentaglycine cross-bridge for cell wall peptidoglycan cross-linking (45) and has previously been linked to phage susceptibility (21).

Beyond cell wall synthesis, clones S1 and K2 contained a substitution in *potA* or truncation in *potB*, respectively. The *potAB* genes are part of the *potABCD* operon shown in *E. coli* to encode a membrane-associated spermidine-preferential uptake system, where *potABC* forms the ABC-transporter (46). Strikingly, three clones (K1, R1, and R3) harbored mutations in the same gene, namely *deoC1*. Indeed, the same *deoC1* substitution (V136L) and truncation (E134*) mutations arose in multiple clones (Table 1). The *deoC1* gene encodes an aldolase, shown in *Salmonella* and *E. coli* to be involved in deoxyribonucleotide catabolism by catalyzing the reversible degradation of 2-deoxy-D-ribose 1-phosphate to D-glyceraldehyde 3-phosphate and acetaldehyde (47). Finally, K3 contained a N-terminal truncation in the *cshA* RNA helicase gene, which has been linked to mRNA stabilization and degradation, including degradation of *S. aureus* mRNAs of the quorum sensing system *agrBDCA* and RNAIII (48).

### CRISPRi validation of identified genes in phage resistance

To confirm that the mutated genes in the selected clones were indeed directly involved in phage resistance, reversible inhibition of gene expression using CRISPR interference (CRISPRi) (39) was utilized in the parental JE2 background. For the different genes, targeted knockdowns were designed using single guide RNA constructs and a vector encoding an IPTG-inducible dead *cas9*. Phage resistance of the gene knockdowns was assessed using liquid growth assays in the absence and presence of low phage concentrations (either MOI 10^-4^ or 10^-5^, as indicated in Fig 2). Increased survival during phage exposure in the IPTG-activated knockdown compared to the uninduced condition was interpreted as increased phage resistance (Fig 2). Gene knockdown led to increased phage resistance for *tagO*, *ligA*, *cshA*, *femA*, *deoC1*, *potAB*, but not for *merR* (Fig 2). This indicates that the genes likely responsible for phage resistance in clones R2 were *ligA* and *femA*, but not *merR*. Gene knockdown in the absence of the phage led to growth defects for *tagO, cshA, and ligA*, and to a lesser extent *femA* and *deoC1*, indicating that these genes are also important for normal bacterial growth.

**Figure 2.**
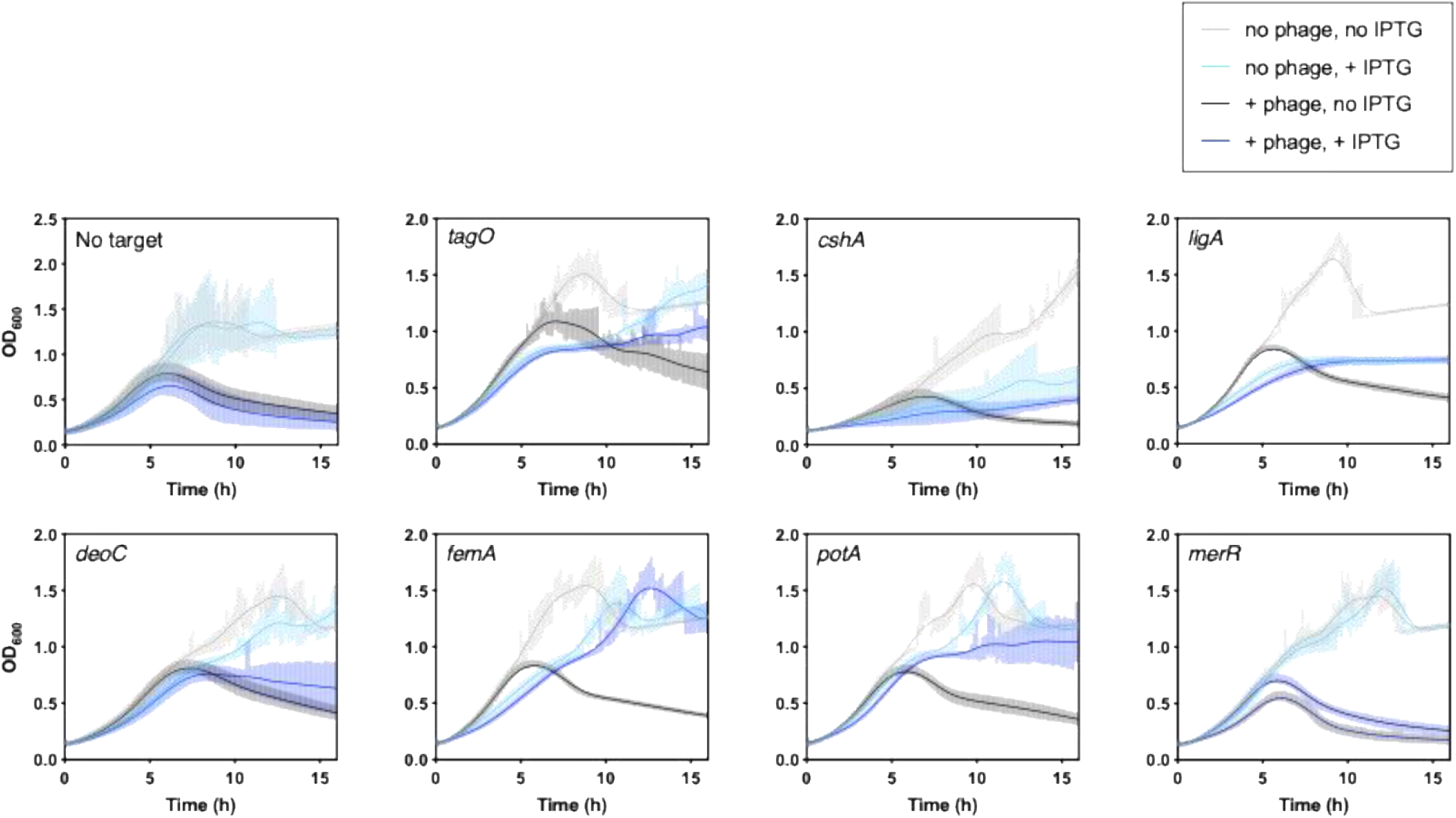
Growth curves of *S. aureus* strains with CRISPRi knockdown of indicated genes in absence and presence of phage. **K.** Graphs show growth curves for *S. aureus* strains with (light/dark blue, 250 µM) or without (grey/black, 0 µM) IPTG-induced CRISPRi knockdown of genes with SNPs in the phage resistant clones. Each graph represents the knockdown targeting one gene, as indicated by the gene name in the top-left corner. As in the legend, graphs plot 4 conditions: exposure to phage buffer only (no phage, no IPTG; grey), with IPTG (no phage, +IPTG: sky blue), with phage K but without IPTG (phage, no IPTG: black), and with phage K and IPTG (phage, +IPTG: dark blue). Phage K concentration was MOI 10^-4^, except for *tagO* and *deoC* where it was MOI^-5^. Graphs show the mean of 6 biological replicates, with shading indicating standard error of mean. The curves representing the mean were smoothed using LOWESS (10 points in smoothing window).

The *deoC1* mutations in the phage resistant clones were all in the same 20 amino acid region located toward the C-terminal, predicted to be within an alpha helix region of the protein. Interestingly, the *deoC1* knockdown only showed a small shift towards better survival compared to uninduced bacteria, suggesting that deletion of *deoC1* only marginally affected phage resistance. However, an N-terminal transposon insertion mutant of *deoC1* showed larger plaque formation and slightly increased phage susceptibility when infected with phage K, supporting that this gene is involved in phage susceptibility (Fig S2G). The limited phage resistance phenotype for the *deoC1* knockdown and reversed effect in the Tn insertion potentially implicates the N-terminal region as necessary for the phage resistance phenotype. However, the *deoC1* data together indicate a novel interaction between phages and the host aldolase, although the mechanism for this remains to be elucidated.

### Phage resistance affects antibiotic susceptibility

Three of the clones had mutations in genes that encode molecules important for cell wall synthesis, specifically *tagO* and *femA*. Consequently, these mutations may produce changes to the cell wall, and with that influencing susceptibility to cell wall-targeting antibiotics (13). Therefore, we determined the minimal inhibitory concentrations (MIC) of the phage-resistant clones for four cell wall-targeting antibiotics: cefoxitin, cefotaxime, oxacillin and vancomycin. The parental JE2 strain was susceptible to vancomycin, but resistant to oxacillin and cefoxitin and intermediate resistant to cefotaxime based on CLSI guidelines (Fig 3 and Supp. F3). For the phage-resistant clones there were changes in the MICs for all antibiotics. Most notably, *cshA* mutant K3 became susceptible to oxacillin (≤2 mg ml^-1^, CLSI guidelines) and cefotaxime (≤8 mg ml^-1^), with 1- to 2-log fold decreases in MIC to these antibiotics (Fig S4). Similarly, and as expected for *S. aureus* with defective TagO, *tagO* mutants S2 and S3 also became susceptible to oxacillin and cefotaxime. However, S2 and S3 showed concomitantly increased MICs towards vancomycin by 1.5-fold to 2 mg ml^-1^. Interestingly, S2 and S3 had similar MICs across all antibiotics, while other clones mutated in the same genes, such as the three *deoC1* aldolase mutants (K1, R1 and R3), showed more variation (Fig 3). For the remaining clones, the differences in MIC were relatively small compared to the parental JE2 strain.

**Figure 3.**
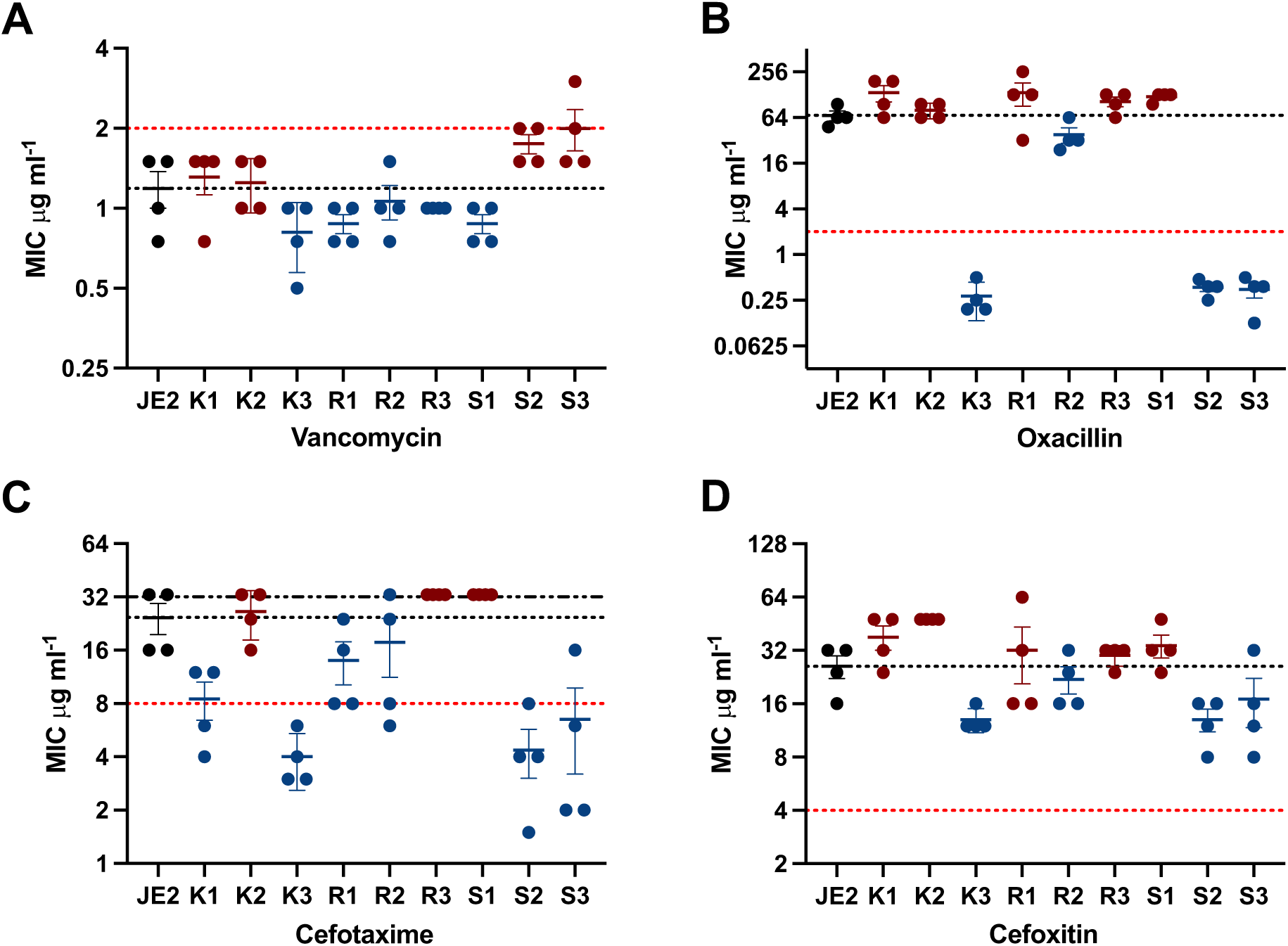
Minimum inhibitory concentration (MIC) of *S. aureus* phage-resistant clones for cell wall targeting antibiotics. Susceptibility of *S. aureus* JE2 wild type and phage-resistant clones towards the indicated antibiotics was determined by E-tests, with breakpoints indicating susceptibility and intermediate susceptible *S. aureus* towards **A** vancomycin, or susceptibility and resistance towards **B** oxacillin, **C** cefotaxime and **D** cefoxitin. Clones with increased susceptibility (blue) and decreased susceptibility (red) are shown compared to JE2 (black dots). CLSI breakpoints are indicated by dotted red lines, the JE2 MIC is marked by a black dotted line, and in **C** upper limits of detection of the E-test are marked by a dashed/dotted black line. Results represent average from four independent biological replicates +/-standard deviation.

We generated a correlation matrix analysis with Pearson correlation coefficients to determine whether phage resistance (PFU ml^-1^) correlated with antibiotic susceptibility (MIC), using Z scores for comparison of the different values (Fig 4). Comparing all strains (Fig 4A), increased resistance to one phage had a strong positive correlation (>0.5) with increased resistance to the other two phages, which may reflect the overlapping pathway of bacterial infection between the phages. Increased phage resistance correlated moderately with decreased β-lactam resistance as indicated by the slight negative correlations (∼-0.5; Fig 4). Analyzing the clones by mutation type, i.e. cell wall (*tagO* and *femA*), aldolase (*deoC1)* or ‘other’ (Figure 4B-D, respectively), revealed that cell wall-related phage resistance (R2, S2, and S3) showed a strong negative correlation between phage resistance and β-lactam resistance (<-0.5, Fig 4B). Conversely, correlations between phage and antibiotic resistance for the aldolase and ‘other’ groupings were much weaker (closer to 0, Fig 4C and D). Overall, our results indicated that phage resistance only correlated strongly with changes in β-lactam antibiotic susceptibility when phage resistance had developed through mutations in genes involved in cell wall biosythesis.

**Figure 4.**
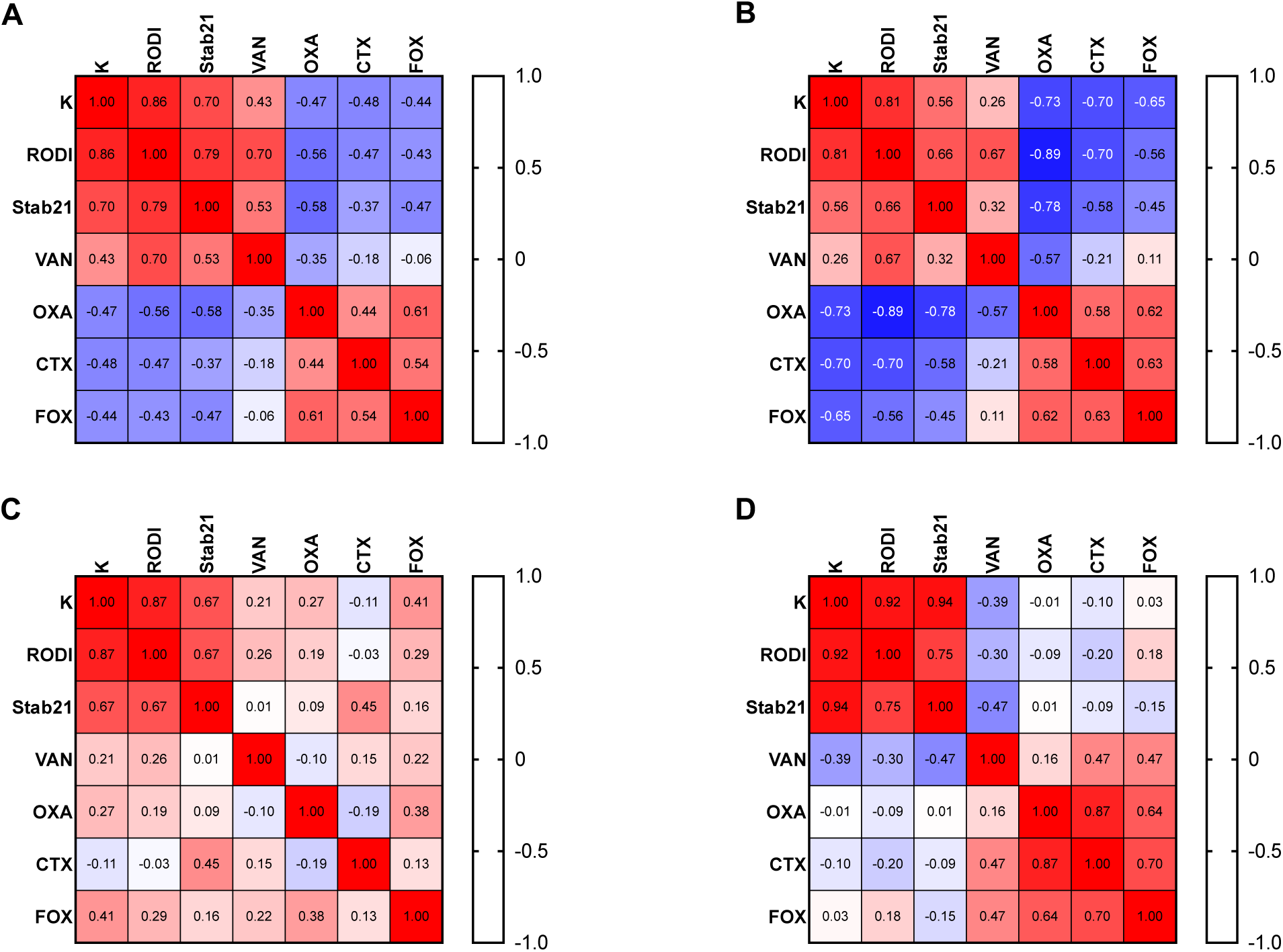
Correlation matrices between phage and antibiotic resistance. Pearson r correlation test with phage PFU ml-1 (K, RODI, Stab21) and E-test MIC results (VAN, OXA, CTX, FOX) for **A.** all phage-resistant clones and parental JE2. **B.** clones with cell wall mutations (R2, S2, S3) and parental JE2. **C.** clones with aldolase mutations (K1, R1, R3) and parental JE2. **D.** for clones grouped as ‘other’ mutations (S1, K2, K3) and parental JE2. In all cases, PFU ml^-1^ were log transformed and inversed to make them align logically with the MIC values (higher values meaning higher resistance), and all values were normalized to a comparable scale using a Z score calculation. Positive and negative correlations are shown in red and blue, respectively.

### Mutations linked to phage resistance lead to altered wall teichoic acid glycosylation

Given the presence of *tagO* and *femA* cell wall mutants, as well as links between phage susceptibility and WTA glycosylation (3, 4, 7), the WTA glycosylation pattern of selected clones was assessed using an antibody deposition assay (Fig 5A-B). α- and β-GlcNAc glycosylation were investigated for the JE2 parental strain, a Δ*tagO* mutant (49) and phage resistant clones with changes in cell wall-related genes *tagO* (S2, S3) and *femA* (R2), which showed increased β-lactam susceptibility. Further, we included clone K3. K3 also showed increased β-lactam susceptibility and is mutated in the *cshA* RNA helicase. The *cshA* RNA helicase mediates *agr* operon and RNAIII mRNA stability, and the *agr* operon has been linked to WTA glycosylation regulation via *tarM* (48). The phage resistant clones with mutations in *tagO* (S2 and S3) had the same glycosylation pattern as the Δ*tagO* mutant, namely undetectable levels of α- and β-GlcNAc (Fig 5B), suggesting that the *tagO* mutations in clones S2 and S3 have disrupted WTA synthesis. R2 containing a *femA* mutation (as well as *ligA* and *merR* mutations) had significantly elevated levels of α-glycosylation compared to the JE2 parental strain. Finally, clone K3 had significantly elevated levels of β-glycosylation. Together, the results show that phage-resistance linked mutations can lead to altered glyco-switching, either towards α- or β-GlcNAc depending on the underlying mutation.

**Figure 5.**
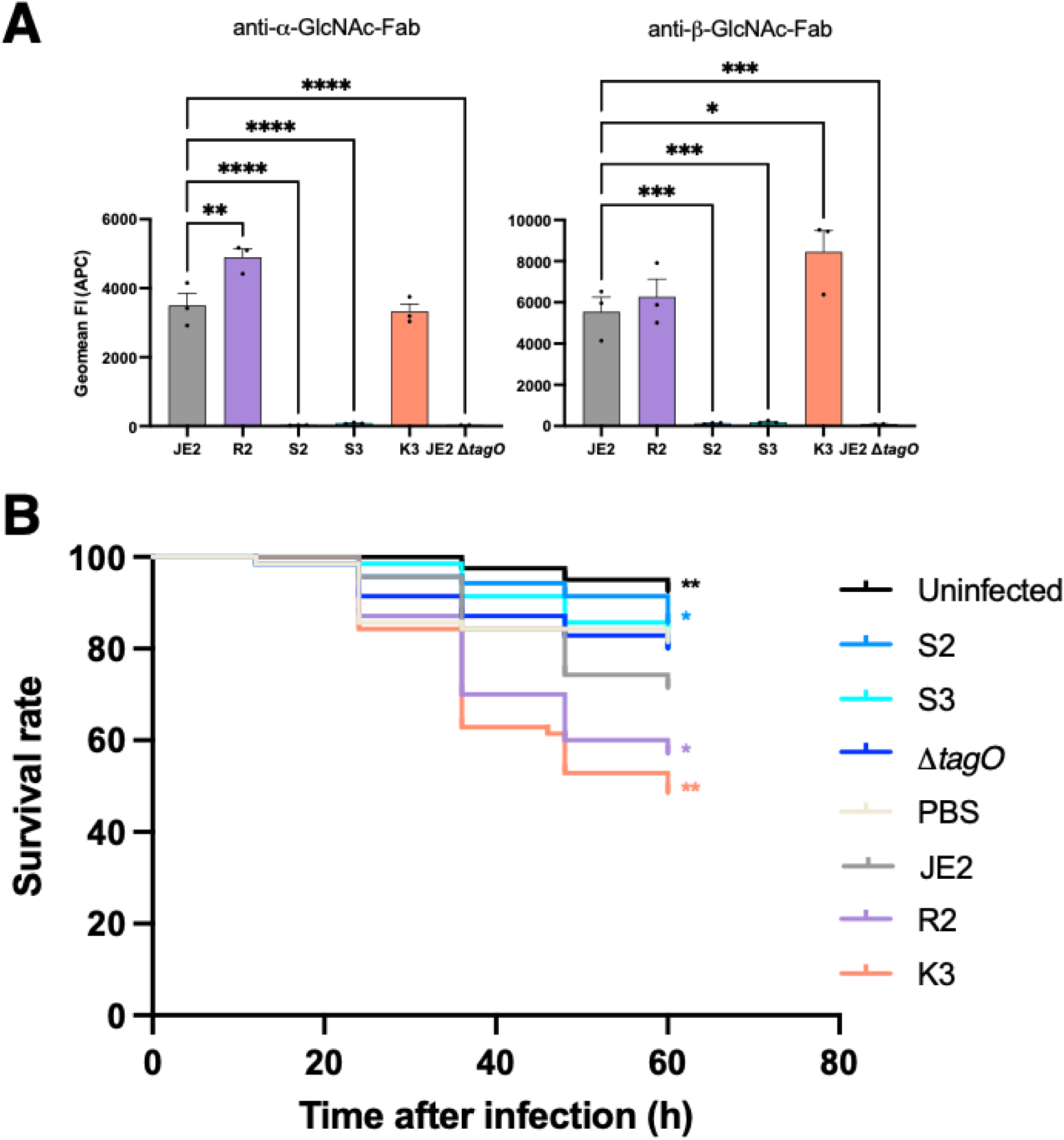
WTA glycosylation pattern and *in vivo* virulence in *Galleria mellonella* of specific phage-resistant clones. **A.** WTA antibody deposition assay for detection of α-1,4- and β-1,4-GlcNAc in JE2 wild-type, indicated clones, and JE2 Δ*tagO* mutant. One-way ANOVA with Dunnett’s correction for multiple comparisons was performed to compare GlcNAc levels of the clones to JE2. **B.** Bacterial virulence depicted by the survival rate of *Galleria mellonella* over 60 h following intrahemocoelic challenge with JE2 parental strain, indicated phage-resistant clones, or JE2 Δ*tagO* mutant. For different strains, means from 7 biological replicates are indicated by line and dot color as shown in legend. Survival statistics were based on the log-rank Mantel-Cox test comparisons between clones and parental JE2. Significant differences are indicated by one (*p* <0.05), two (*p* <0.01), three (*p* <0.001) or four (*p* <0.0001) asterisks (*).

### The phage resistant clones displayed altered virulence in a Galleria *in vivo* model

In *S. aureus*, WTA glycosylation mediates specific immune interactions with the host (5). Indeed, expression and predominance of either α- or β-glycosylation is affected by environmental conditions, where β-glycosylation is upregulated *in vivo* (50). Given the altered WTA glycosylation phenotypes for R2 and K3 and disrupted phenotype of S2 and S3, we compared the virulence of these strains to JE2 and *ΔtagO*, using our well-established *Galleria mellonella* infection model. The *tagO* mutant clones S2 and S3 behaved similarly to *ΔtagO*, with S2 showing significantly reduced killing of *Galleria mellonella* compared to JE2 wildtype (Figure 5C). On the other hand, R2 and K3 resulted in increased killing of *Galleria mellonella* in comparison to the wildtype (Figure 5C). For *cshA* mutant K3, this may be associated with the increased levels of β-GlcNAc (Figure 5B), which has previously been linked to immune stimulatory effects (51). Further, mutations in *cshA* have been shown to stabilize *agr* mRNA, the quorum sensing system that regulates many virulence factors (48). It is unclear what is responsible for the increased virulence of R2, with increased α-glycosylation compared to the parental strain and multiple genes affected by mutations (*femA*, *ligA*, *merR*). Overall, the *Galleria mellonella* model data supports previous findings that *tagO* mutants have attenuated virulence. It is interesting that R2 and K3, with differentially altered WTA glycosylation, are both more virulent than the parental strain. Potentially, virulence could be enhanced through other routes than WTA glycosylation, such as changed virulence factor expression. Likewise, the glycosylation pattern of the clones could differ in the *Galleria mellonella* model compared to the *in vitro* conditions of the antibody disposition assay, particularly given that β-GlcNAc switching has been shown to occur during *in vivo* conditions (50).

## Discussion

Our study showed that reduced phage susceptibility in several clones of *S. aureus* was associated with mutations in *femA* and *tagO*, two genes linked to cell wall biosynthesis. This finding confirms findings from previous studies where mutations in genes linked to WTA biosynthesis (including *tagO* and teichoic acid ribitol (Tar) enzymes *tarA*, *tarK*) (52) and in *femA* were identified (20, 21). TarA catalyzes the synthesis of the intermediate lipid β [ManNAc-β-(1-4)-GlcNAc-P-P-C55], while TarK is involved in RboP priming and polymerization (6, 53). Deletion of *tagO* or *tarA* in *S. aureus* abolishes WTA biosynthesis (54), while bacteria with a *tarK* deletion produce longer, more heterogeneous WTA (53). The exact mechanism of how *femA* mutations lead to phage resistance is less clear. FemA catalyzes the formation of pentaglycine cross-bridges for peptidoglycan cross-linking. Deletion of *femA* has been shown to lead to shortened cross-bridge formation, reduced peptidoglycan cross-linking, partial cell-wall thickening, and accumulation of immature peptidoglycan at the cell septum, suggesting a changed cell-wall architecture (55). As phages have to enzymatically degrade the peptidoglycan by cell wall hydrolases (56), to enter bacterial cells, it is likely that changes in the architecture of the peptidoglycan could interfere with phage entry.

We also observed that mutations in *potA* and *potB* resulted in decreased phage susceptibility. The *potABCD* operon forms a membrane-bound ABC transporter, where PotABC comprises the transporter and PotD binds the substrate (46). PotABCD is involved in the entry of polyamines into the cell (46). Interestingly, polyamines have been shown to both facilitate efficient phage DNA packaging in phage heads (57) but also to inhibit phage injection and replication at high levels (58, 59). Further, a study utilized *potABCD* transposon and knockout mutants as phage insensitive strain backgrounds (60), again linking the system to phage susceptibility.

Several of the phage resistant clones evolved in this study had mutations in genes not previously linked to phage resistance. Mutated in K3, the *cshA* DEAD box RNA helicase of *S. aureus* regulates the stabilization and degradation of numerous mRNAs, including regulatory protein *sarA* mRNA (61), *agr* quorum sensing operon mRNA (48) and mRNA from the pyruvate dehydrogenase (PDH) operon (62). Interestingly, accumulation of *pdh* mRNA in *cshA* mutants exacerbated the production of straight-chain fatty acids, and such shifts in fatty acid metabolism can lead to decreased membrane fluidity (63). Reduced membrane fluidity has been linked to phage resistance in *Bacillus subtilus*, where stabilization of the membrane by bacterial dynamin-like protein DynA had a protective effect against cell lysis and phage progeny release (64). DNA ligase A (*ligA*), one of three genes mutated in R2, is an essential, NAD^+^-dependent enzyme involved in DNA replication and repair by catalyzing phosphodiester linkage formation between 5’-phosphoryl and 3’-hydroxyl groups (43). Phages have been shown to encode their own ligase genes (28, 65), while they also utilize the host ligase for gap sealing during both theta and rolling circle phage DNA replication (66). Furthermore, *ligA* was downregulated in *Lactococcus lactis* during phage infection (67). Interestingly, the CRISPRi screen performed in this study indicates that *ligA* and *cshA* are required for normal growth in laboratory media conditions without phage challenge (Fig 2). The *deoC1* gene was mutated in three separate clones, K1, R1, and R3. DeoC1 catalyzes the reversible degradation of 2-deoxy-D-ribose 1-phosphate to D-glyceraldehyde 3-phosphate and acetaldehyde (47). It has been shown that *deoC1* is induced by the presence of external DNA (68) and it has been suggested it may prevent uptake of foreign DNA, albeit without a defined mechanism (69). Likewise, *deoC* was downregulated in *Bacillus subtilis* undergoing lytic phage infection (70).

The relationship between phage- and antibiotic resistance is increasingly of interest as phage-antibiotic combination treatments are proposed as a potential solution to treating infections with antimicrobial resistant bacteria. While some research indicates an evolutionary trade-off between phage and antibiotic resistance, other results suggest a more antagonistic relationship where phage resistance may also confer antibiotic resistance (71). In this study, we did not identify a general link between *S. aureus* phage and antibiotic resistance. In line with previous findings (4), we observed a strong correlation between phage resistance and increased β-lactam susceptibility, but mostly for clones containing mutations in genes implicated in cell wall biosynthesis (S2, S3, and R2; *tagO* or *femA*). When clones evolved phage resistance via mutations in other pathways, there was no increase in β-lactam susceptibility. One exception to this was K3, containing mutations in *cshA*, which even became susceptible to β-lactams. Here, it is likely that combined changes to *sarA*, *agr* and *pdh* mRNA stabilization could have downstream effects on the cell wall and membrane.

In contrast to *E. coli*, where numerous outer membrane phage receptors have been identified (72), *S. aureus* phages characterized so far all recognize epitopes of WTA as their primary receptor. In *S. aureus*, WTA is associated with host colonization and is a key site for interaction between the bacterial cell and the host immune system (5). WTA expression and upregulation of β-glycosylation has been linked to increased strain prevalence *in vivo* (50), and an absence of WTA reduces the colonization and virulence capability of *S. aureus*. Indeed, WTA-deficient *ΔtagO* mutants had impaired adherence to human epithelial (24) and endothelial cells (23), and showed attenuated colonization and virulence in *in vivo* models (23). We confirmed that the phage-resistant clones with mutations in *tagO* evolved in this study lacked WTA expression, since we could not detect α- or β-GlcNAc expression and observed similar attenuation of growth, hemolysis and virulence compared to the *ΔtagO* mutant. Thus, it is an ongoing question whether deletion or alteration of the WTA phage receptor would be a viable route of *S. aureus* phage resistance *in vivo*.

Taken together, our results implicate several novel bacterial pathways in phage resistance. Furthermore, we find that relationships between phage resistance, antibiotic susceptibility and virulence are dependent on the pathways and mutations leading to phage resistance.

### Opening up

This work gives an insight into the complexity of phage-bacteria interactions. Several novel mutations linked to the development of phage resistance are identified, including *deoC1* aldolase, *ligA* DNA ligase, *cshA* RNA helicase and *potAB* ABC putrescine transporter mutations. Potentially, phage resistance from these mutations could occur by interference during the phage lytic cycle following phage DNA injection into the bacterial cell, although downstream effects on the cell wall could be possible. Next steps will be to elucidate the mechanisms underlying the phage-host interactions related to these genes, and how the mutations identified here confer phage resistance. This work differs to previous studies on phage resistance in *S. aureus*, where most mutations were found in genes associated with WTA and peptidoglycan synthesis (e.g. *tagO, tarA/K,* and *femA*). In these cases, resistance likely occurs via depletion or alteration of the phage receptor, or interference with the entry of the phage into the bacterial cell. Although this study also identified phage resistant clones with mutations in WTA and cell wall synthesis genes (*tagO* and *femA*), these made up only one third of the total clones. Here, the WTA *tagO* mutants showed reduced virulence in an *in vivo* G*alleria mellonella* model. Indeed, cell wall mutants, particularly those depleted in WTA, often show aberrant growth *in vitro* and fail to colonize the host *in vivo*. Thus, there is a question on their clinical relevance. Interestingly, in the largest published analysis of phage therapy cases to date, no phage resistance was observed for the limited number of *S. aureus* samples obtained from patients undergoing phage therapy for *S. aureus* infections (1). In the current study, clones mutated in *cshA* and *ligA*/*femA* had increased virulence in a *Galleria mellonella* model. Thus, it could be relevant to test their potential for viability in an *in vivo* or humanized setting.

## Acknowledgements

Thanks to the Master’s students on the CRISPR Tsunami course for testing our CRISPRi phage susceptibility method. Thanks to Rob van Dalen for technical assistance with the WTA antibody assays. Thank you to Pilar Garcia for providing us with the ϕIPLA-RODI phage. Thank you to Mikael Skurnik for providing us with the Stab21 phage. Thank you to Janes Kruse for providing us with phage K. Thanks to Zhian Salehian for help with cloning.

## CRediT authorship contribution statement

**Janine Bowring:** Writing – review & editing, Writing – original draft, Visualization, Methodology, Investigation, Formal analysis, Conceptualization, Supervision, Project administration **Freja C. Mikkelsen:** Writing – review & editing, Writing – original draft, Visualization, Methodology, Investigation, Formal analysis **Roshni Haider:** Methodology, Investigation, Formal analysis **Esther Lehmann:** Investigation, Formal analysis, Writing – review & editing **Thibault Frisch:** Methodology, Investigation, Formal analysis, Writing-review **Morten Kjos:** Resources, Methodology, Writing-review and editing **Nina van Sorge:** Resources, Supervision, Writing-review and editing **Hanne Ingmer:** Conceptualization, Supervision, Project administration, Funding acquisition, Writing – review & editing

## References

1. Naghavi M, Vollset SE, Ikuta KS, Swetschinski LR, Gray AP, Wool EE, et al. Global burden of bacterial antimicrobial resistance 1990–2021: a systematic analysis with forecasts to 2050. The Lancet. 2024;404(10459):1199–226.

2. Pirnay J-P, Djebara S, Steurs G, Griselain J, Cochez C, De Soir S, et al. Personalized bacteriophage therapy outcomes for 100 consecutive cases: a multicentre, multinational, retrospective observational study. Nature Microbiology. 2024;9(6):1434–53.

3. Ingmer H, Gerlach D, Wolz C. Temperate Phages of Staphylococcus aureus. Microbiology Spectrum. 2019;7(5).

4. Krusche J, Beck C, Lehmann E, Gerlach D, Daiber E, Mayer C, et al. Characterization and host range prediction of Staphylococcus aureus phages through receptor-binding protein analysis. Cell Reports. 2025;44(3):115369.

5. van Dalen R, Peschel A, Van Sorge NM. Wall Teichoic Acid in Staphylococcus aureus Host Interaction. Trends in Microbiology. 2020;28(12):985–98.

6. Weidenmaier C, Lee JC. Structure and Function of Surface Polysaccharides of Staphylococcus aureus. Current Topics in Microbiology and Immunology: Springer International Publishing; 2015. p. 57–93.

7. Yang J, Bowring JZ, Krusche J, Lehmann E, Bejder BS, Silva SF, et al. Cross-species communication via agr controls phage susceptibility in Staphylococcus aureus. Cell Reports. 2023;42(9):113154.

8. Kuijk MM, Tusveld E, Lehmann E, van Dalen R, Lasa I, Ingmer H, et al. The two-component system ArlRS is essential for wall teichoic acid glycoswitching in Staphylococcus aureus. mBio. 2025;16(1):e0266824.

9. Xu X, Gu P. Overview of Phage Defense Systems in Bacteria and Their Applications. International Journal of Molecular Sciences. 2024;25(24):13316.

10. Mikkelsen K, Bowring JZ, Ng YK, Svanberg Frisinger F, Maglegaard JK, Li Q, et al. An Endogenous Staphylococcus aureus CRISPR-Cas System Limits Phage Proliferation and Is Efficiently Excised from the Genome as Part of the SCCmec Cassette. Microbiol Spectr. 2023;11(4):e0127723.

11. Kim MK, Chen Q, Echterhof A, Pennetzdorfer N, Mcbride RC, Banaei N, et al. A blueprint for broadly effective bacteriophage-antibiotic cocktails against bacterial infections. Nature Communications. 2024;15(1).

12. Müller DM, Pourtois JD, Kim MK, Targ B, Burgener EB, Milla C, et al. Bacterial Receptors but Not Anti-Phage Defence Mechanisms Determine Host Range for a Pair of Pseudomonas aeruginosa Lytic Phages. 2024.

13. Farha MA, Leung A, Sewell EW, D’Elia MA, Allison SE, Ejim L, et al. Inhibition of WTA synthesis blocks the cooperative action of PBPs and sensitizes MRSA to beta-lactams. ACS Chem Biol. 2013;8(1):226–33.

14. Sobhanifar S, Worrall LJ, King DT, Wasney GA, Baumann L, Gale RT, et al. Structure and Mechanism of Staphylococcus aureus TarS, the Wall Teichoic Acid β-glycosyltransferase Involved in Methicillin Resistance. PLOS Pathogens. 2016;12(12):e1006067.

15. Bertsche U, Weidenmaier C, Kuehner D, Yang SJ, Baur S, Wanner S, et al. Correlation of daptomycin resistance in a clinical Staphylococcus aureus strain with increased cell wall teichoic acid production and D-alanylation. Antimicrob Agents Chemother. 2011;55(8):3922–8.

16. Hort M, Bertsche U, Nozinovic S, Dietrich A, Schrötter AS, Mildenberger L, et al. The Role of β-Glycosylated Wall Teichoic Acids in the Reduction of Vancomycin Susceptibility in Vancomycin-Intermediate Staphylococcus aureus. Microbiology Spectrum. 2021;9(2).

17. Jo A, Ding T, Ahn J. Synergistic antimicrobial activity of bacteriophages and antibiotics against Staphylococcus aureus. Food Science and Biotechnology. 2016;25(3):935–40.

18. Mehmood Z, Kanwar R, Ullah K, Ali S, Aslam MA, Qadeer A, et al. Synergistic effects of zP-1 phage and ampicillin against methicillin-resistant Staphylococcus aureus isolated from hospital staff. Irish Journal of Medical Science (1971 -). 2025;194(2):611–21.

19. Simon K, Pier W, Krüttgen A, Horz H-P. Synergy between Phage Sb-1 and Oxacillin against Methicillin-Resistant Staphylococcus aureus. Antibiotics. 2021;10(7):849.

20. Tran M, Hernandez Viera AJ, Tran PQ, Mo CY. Bacteriophage infection drives loss of β-lactam resistance in methicillin-resistant Staphylococcus aureus. eLife. 2024.

21. Berryhill BA, Huseby DL, McCall IC, Hughes D, Levin BR. Evaluating the potential efficacy and limitations of a phage for joint antibiotic and phage therapy of Staphylococcus aureus infections. Proc Natl Acad Sci U S A. 2021;118(10):e2008007118.

22. Jung D-J, An J-H, Kurokawa K, Jung Y-C, Kim M-J, Aoyagi Y, et al. Specific Serum Ig Recognizing Staphylococcal Wall Teichoic Acid Induces Complement-Mediated Opsonophagocytosis against Staphylococcus aureus. The Journal of Immunology. 2012;189(10):4951–9.

23. Weidenmaier C, Peschel A, Xiong YQ, Kristian SA, Dietz K, Yeaman MR, et al. Lack of wall teichoic acids in Staphylococcus aureus leads to reduced interactions with endothelial cells and to attenuated virulence in a rabbit model of endocarditis. J Infect Dis. 2005;191(10):1771–7.

24. Baur S, Rautenberg M, Faulstich M, Grau T, Severin Y, Unger C, et al. A Nasal Epithelial Receptor for Staphylococcus aureus WTA Governs Adhesion to Epithelial Cells and Modulates Nasal Colonization. PLoS Pathogens. 2014;10(5):e1004089.

25. Weidenmaier C, Mcloughlin RM, Lee JC. The Zwitterionic Cell Wall Teichoic Acid of Staphylococcus aureus Provokes Skin Abscesses in Mice by a Novel CD4+ T-Cell-Dependent Mechanism. PLoS ONE. 2010;5(10):e13227.

26. Brignoli T, Douglas E, Duggan S, Fagunloye OG, Adhikari R, Aman MJ, et al. Wall Teichoic Acids Facilitate the Release of Toxins from the Surface of Staphylococcus aureus. Microbiology Spectrum. 2022;10(4).

27. Wanner S, Schade J, Keinhörster D, Weller N, George SE, Kull L, et al. Wall teichoic acids mediate increased virulence in Staphylococcus aureus. Nature Microbiology. 2017;2(4):16257.

28. O’Flaherty S, Coffey A, Edwards R, Meaney W, Fitzgerald GF, Ross RP. Genome of staphylococcal phage K: a new lineage of Myoviridae infecting gram-positive bacteria with a low G+C content. J Bacteriol. 2004;186(9):2862–71.

29. Gutiérrez D, Vandenheuvel D, Martínez B, Rodríguez A, Lavigne R, García P. Two Phages, phiIPLA-RODI and phiIPLA-C1C, Lyse Mono- and Dual-Species Staphylococcal Biofilms. Applied and Environmental Microbiology. 2015;81(10):3336–48.

30. Oduor JMO, Kadija E, Nyachieo A, Mureithi MW, Skurnik M. Bioprospecting Staphylococcus Phages with Therapeutic and Bio-Control Potential. Viruses. 2020;12(2):133.

31. Krusche J, Beck C, Lehmann E, Gerlach D, Daiber E, Mayer C, et al. Characterization and host range prediction of Staphylococcus aureus phages through receptor-binding protein analysis. Cell Rep. 2025;44(3):115369.

32. Bowring JZ, Su Y, Alsaadi A, Svenningsen SL, Parkhill J, Ingmer H. Screening for Highly Transduced Genes in Staphylococcus aureus Revealed Both Lateral and Specialized Transduction. Microbiology Spectrum. 2022;10(1).

33. Virtanen P, Gommers R, Oliphant TE, Haberland M, Reddy T, Cournapeau D, et al. SciPy 1.0: fundamental algorithms for scientific computing in Python. Nature Methods. 2020;17(3):261–72.

34. Cheung GYC, Duong AC, Otto M. Direct and synergistic hemolysis caused by Staphylococcus phenol-soluble modulins: implications for diagnosis and pathogenesis. Microbes and Infection. 2012;14(4):380–6.

35. Adhikari RP, Arvidson S, Novick RP. A nonsense mutation in agrA accounts for the defect in agr expression and the avirulence of Staphylococcus aureus 8325-4 traP::kan. Infect Immun. 2007;75(9):4534–40.

36. Hendriks A, van Dalen R, Ali S, Gerlach D, van der Marel GA, Fuchsberger FF, et al. Impact of Glycan Linkage to Staphylococcus aureus Wall Teichoic Acid on Langerin Recognition and Langerhans Cell Activation. ACS Infect Dis. 2021;7(3):624–35.

37. Driguez P-A, Guillo N, Rokbi B, Mistretta N, Talaga P, inventors; Sanofi Pasteur, assignee. Immunogenic Compositions Against S. aureus. WO patent WO 2017/064190 A1. 2017 2016/10/13.

38. Sprouffske K, Wagner A. Growthcurver: an R package for obtaining interpretable metrics from microbial growth curves. BMC Bioinformatics. 2016;17(1).

39. Liu X, de Bakker V, Heggenhougen MV, Marli MT, Froynes AH, Salehian Z, et al. Genome-wide CRISPRi screens for high-throughput fitness quantification and identification of determinants for dalbavancin susceptibility in Staphylococcus aureus. mSystems. 2024;9(7):e0128923.

40. Galperin MY, Alvarez V, Roberto, Karamycheva S, Makarova KS, Wolf I, Yuri, Landsman D, et al. COG database update 2024. Nucleic Acids Research. 2025;53(D1):D356–D63.

41. Mistry J, Chuguransky S, Williams L, Qureshi M, Salazar A, Gustavo, Sonnhammer ELL, et al. Pfam: The protein families database in 2021. Nucleic Acids Research. 2021;49(D1):D412–D9.

42. Jurado A, Fernández L, Rodríguez A, García P. Understanding the Mechanisms That Drive Phage Resistance in Staphylococci to Prevent Phage Therapy Failure. Viruses. 2022;14(5):1061.

43. Kaczmarek FS, Zaniewski RP, Gootz TD, Danley DE, Mansour MN, Griffor M, et al. Cloning and functional characterization of an NAD(+)-dependent DNA ligase from Staphylococcus aureus. J Bacteriol. 2001;183(10):3016–24.

44. Brown NL, Stoyanov JV, Kidd SP, Hobman JL. The MerR family of transcriptional regulators. FEMS Microbiol Rev. 2003;27(2-3):145–63.

45. Schneider T, Senn MM, Berger-Bachi B, Tossi A, Sahl HG, Wiedemann I. In vitro assembly of a complete, pentaglycine interpeptide bridge containing cell wall precursor (lipid II-Gly5) of Staphylococcus aureus. Mol Microbiol. 2004;53(2):675–85.

46. Qiao Z, Do PH, Yeo JY, Ero R, Li Z, Zhan L, et al. Structural insights into polyamine spermidine uptake by the ABC transporter PotD-PotABC. Science Advances. 2024;10(38).

47. Hoffee PA. 2-deoxyribose-5-phosphate aldolase of Salmonella typhimurium: purification and properties. Arch Biochem Biophys. 1968;126(3):795–802.

48. Oun S, Redder P, Didier JP, Francois P, Corvaglia AR, Buttazzoni E, et al. The CshA DEAD-box RNA helicase is important for quorum sensing control in Staphylococcus aureus. RNA Biol. 2013;10(1):157–65.

49. Slavetinsky J, Lehmann E, Slavetinsky C, Gritsch L, van Dalen R, Kretschmer D, et al. Wall Teichoic Acid Mediates Staphylococcus aureus Binding to Endothelial Cells via the Scavenger Receptor LOX-1. ACS Infectious Diseases. 2023;9(11):2133–40.

50. Winstel V, Kühner P, Salomon F, Larsen J, Skov R, Hoffmann W, et al. Wall Teichoic Acid Glycosylation Governs Staphylococcus aureus Nasal Colonization. mBio. 2015;6(4):e00632–15.

51. Gerlach D, Guo Y, De Castro C, Kim S-H, Schlatterer K, Xu F-F, et al. Methicillin-resistant Staphylococcus aureus alters cell wall glycosylation to evade immunity. Nature. 2018;563(7733):705-9.

52. Royet K, Blazere L, Helluin E, Coignet L, Plumet L, Bonhomme M, et al. Exploring Phage-Staphylococcus aureus Host Dynamics Through Innovative In Vitro Experiment and Pharmacokinetic/pharmacodynamic Modeling. 2025.

53. Meredith TC, Swoboda JG, Walker S. Late-stage polyribitol phosphate wall teichoic acid biosynthesis in Staphylococcus aureus. Journal of Bacteriology. 2008;190(8):3046–56.

54. D’Elia MA, Henderson JA, Beveridge TJ, Heinrichs DE, Brown ED. The N-acetylmannosamine transferase catalyzes the first committed step of teichoic acid assembly in Bacillus subtilis and Staphylococcus aureus. J Bacteriol. 2009;191(12):4030–4.

55. Sharif S, Kim SJ, Labischinski H, Schaefer J. Characterization of Peptidoglycan in Fem-Deletion Mutants of Methicillin-Resistant Staphylococcus aureus by Solid-State NMR. Biochemistry. 2009;48(14):3100–8.

56. Kizziah JL, Manning KA, Dearborn AD, Dokland T. Structure of the host cell recognition and penetration machinery of a Staphylococcus aureus bacteriophage. PLOS Pathogens. 2020;16(2):e1008314.

57. Syvanen M, Yin J. Studies of DNA packaging into the heads of bacteriophage lambda. J Mol Biol. 1978;126(3):333–46.

58. Harrison DP, Bode VC. Putrescine and certain polyamines can inhibit DNA injection from bacteriophage lambda. J Mol Biol. 1975;96(3):461–70.

59. De Mattos CD, Faith DR, Nemudryi AA, Schmidt AK, Bublitz DC, Hammond L, et al. Polyamines and linear DNA mediate bacterial threat assessment of bacteriophage infection. Proceedings of the National Academy of Sciences. 2023;120(9).

60. Lehman SM, Kongari R, Glass AM, Koert M, Ray MD, Plaut RD, et al. Phage K gp102 Drives Temperature-Sensitive Antibacterial Activity on USA300 MRSA. Viruses. 2022;15(1):17.

61. Kim S, Corvaglia A-R, Léo S, Cheung A, Francois P. Characterization of RNA Helicase CshA and Its Role in Protecting mRNAs and Small RNAs of Staphylococcus aureus Strain Newman. Infection and Immunity. 2016;84(3):833–44.

62. Khemici V, Prados J, Petrignani B, Di Nolfi B, Bergé E, Manzano C, et al. The DEAD-box RNA helicase CshA is required for fatty acid homeostasis in Staphylococcus aureus. PLOS Genetics. 2020;16(7):e1008779.

63. Sen S, Sirobhushanam S, Johnson SR, Song Y, Tefft R, Gatto C, et al. Growth-Environment Dependent Modulation of Staphylococcus aureus Branched-Chain to Straight-Chain Fatty Acid Ratio and Incorporation of Unsaturated Fatty Acids. PLOS ONE. 2016;11(10):e0165300.

64. Guo L, Sattler L, Shafqat S, Graumann PL, Bramkamp M. A Bacterial Dynamin-Like Protein Confers a Novel Phage Resistance Strategy on the Population Level in Bacillus subtilis. mBio. 2022;13(1).

65. Wang J, Liu F, Su T, Chang Y, Guo Q, Wang Q, et al. The phage T4 DNA ligase in vivo improves the survival-coupled bacterial mutagenesis. Microbial Cell Factories. 2019;18(1).

66. Weigel C, Seitz H, Abril AM, Salas M, Andreu JM, Hermoso JM, et al. Bacteriophage replication modules. FEMS microbiology reviews. 2006;30(3):321–81.

67. Fallico V, Ross RP, Fitzgerald GF, Mcauliffe O. Genetic Response to Bacteriophage Infection in Lactococcus lactis Reveals a Four-Strand Approach Involving Induction of Membrane Stress Proteins, Alanylation of the Cell Wall, Maintenance of Proton Motive Force, and Energy Conservation. Journal of Virology. 2011;85(22):12032–42.

68. Sgarrella F, Poddie FP, Meloni MA, Sciola L, Pippia P, Tozzi MG. Channelling of deoxyribose moiety of exogenous DNA into carbohydrate metabolism: role of deoxyriboaldolase. Comp Biochem Physiol B Biochem Mol Biol. 1997;117(2):253–7.

69. Hammer-Jespersen K. Nucleoside catabolism. In: Munch-Petersen A, editor. Metabolism of nucleotides, nucleosides and nucleobases in microorganisms. Copenhagen, Denmark: Academic Press; 1983. p. 203-58.

70. Mojardín L, Salas M. Global Transcriptional Analysis of Virus-Host Interactions between Phage ϕ29 and Bacillus subtilis. Journal of Virology. 2016;90(20):9293–304.

71. Kunz Coyne AJ, Eshaya M, Bleick C, Vader S, Biswas B, Wilson M, et al. Exploring synergistic and antagonistic interactions in phage-antibiotic combinations against ESKAPE pathogens. Microbiology Spectrum. 2024;12(10).

72. Hantke K. Compilation of Escherichia coli K-12 outer membrane phage receptors – their function and some historical remarks. FEMS Microbiology Letters. 2020;367(2).

