## Supplementary Files for "Phage resistance impacts antibiotic susceptibility and virulence in *Staphylococcus aureus*"

**Table S1. Bacterial strains used in this study**

| Bacteria | Strain no. | Strain description | Source |
| --- | --- | --- | --- |
| *S. xylosus* | HI5917 | DD-34, propagative host for Stab21 | *(1)* |
| *S. aureus* | HI5882 | USA300 JE2 | *(2)* |
|  | HI5830 | USA300 JE2 Δ*tagO* | *(3)* |
|  | HI2157 | RN4220, phage cured lab strain | *(4)* |
|  | HI2156 | 8325-4, 8325 with prophages ɸ11, ɸ12, and ɸ13 removed, propagative host for ɸIPLA-RODI and K | *(5)* |
|  | HI5911 | K4 (RH7), phage resistant JE2 mutant | *This study* |
|  | HI5912 | R4 (RH9), phage resistant JE2 mutant | *This study* |
|  | HI5881 | K1 (RH10), phage resistant JE2 mutant | *This study* |
|  | HI5913 | S4 (RH11), phage resistant JE2 mutant | *This study* |
|  | HI5882 | R1 (RH12), phage resistant JE2 mutant | *This study* |
|  | HI5883 | R2 (RH18), phage resistant JE2 mutant | *This study* |
|  | HI5914 | K5 (RH19), phage resistant JE2 mutant | *This study* |
|  | HI5884 | S1 (RH20), phage resistant JE2 mutant | *This study* |
|  | HI5915 | R5 (RH21), phage resistant JE2 mutant | *This study* |
|  | HI5885 | K2 (RH22), phage resistant JE2 mutant | *This study* |
|  | HI5886 | S2 (RH23), phage resistant JE2 mutant | *This study* |
|  | HI5887 | S3 (RH30), phage resistant JE2 mutant | *This study* |
|  | HI5888 | K3 (RH31), phage resistant JE2 mutant | *This study* |
|  | HI5889 | R3 (RH32), phage resistant JE2 mutant | *This study* |
|  | HI5614 | JE2 pLOW-dcas9_aad9 | *(6)* |
|  | HI5620 | JE2 pLOW-dcas9_aad9, pVL2336-no target | *This study* |
|  | HI5916 | JE2 pLOW-dcas9_aad9, pVL2336-tagO | *This study* |
|  | HI5890 | JE2 pLOW-dcas9_aad9, pVL2336-cshA | *This study* |
|  | HI5895 | JE2 pLOW-dcas9_aad9, pVL2336-ligA | *This study* |
|  | HI5891 | JE2 pLOW-dcas9_aad9, pVL2336-deoC1 | *This study* |
|  | HI5896 | JE2 pLOW-dcas9_aad9, pVL2336-femA | *This study* |
|  | HI5894 | JE2 pLOW-dcas9_aad9, pVL2336-potAB | *This study* |
|  | HI5892 | JE2 pLOW-dcas9_aad9, pVL2336-merR | *This study* |
| *E. coli* | HI4273 | IM08B | *(7)* |
|  | HI5640 | IM08B pLOW-dcas9_aad9 | *(8)* |
|  | HI5641 | IM08B pVL2336-tagO | *(9)* |
|  | HI5875 | IM08B pVL2336-cshA | *This study* |
|  | HI5878 | IM08B pVL2336-ligA | *This study* |
|  | HI5873 | IM08B pVL2336-deoC1 | *This study* |
|  | HI5874 | IM08B pVL2336-femA | *This study* |
|  | HI5876 | IM08B pVL2336-potAB | *This study* |
|  | HI5877 | IM08B pVL2336-merR | *This study* |

**Table S2. Lytic bacteriophages used in this study.**

| Bacteriophages | Propagative strain | Source |
| --- | --- | --- |
| ɸIPLA-RODI | *S. aureus* 8325-4 | *(10)* |
| K | *S. aureus* 8325-4 | *(11)* |
| Stab21 | *S. xylosus* DD-34 | *(1)* |

**Table S3. Antibody-related reagents and resources.**

| **Name** | **SOURCE** | **IDENTIFIER** |
| --- | --- | --- |
| Clone 4461: anti-α-1,4-GlcNAc-WTA Fab | Nina van Sorge Amsterdam UMC, NL (12) | N/A |
| Clone 4462: anti-β-1,4-GlcNAc-WTA Fab | Nina van Sorge Amsterdam UMC, NL (12) | N/A |
| Goat F(ab')2 anti-human-kappa-AF647, secondary antibody | SouthernBiotech | 2062-31 |
| BSA (bovine serum albumin) | Sigma-Aldrich | Cat#A7030 |

**Table S4. Gene putative function from COG/PFAM associations.**

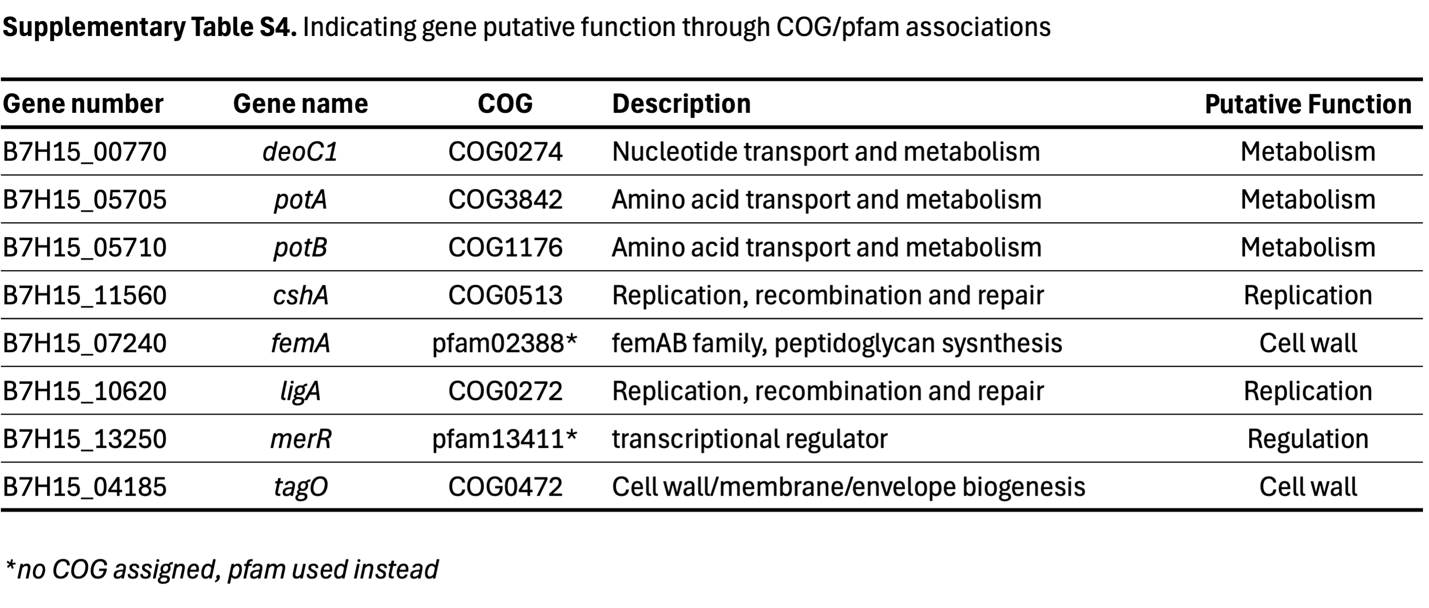

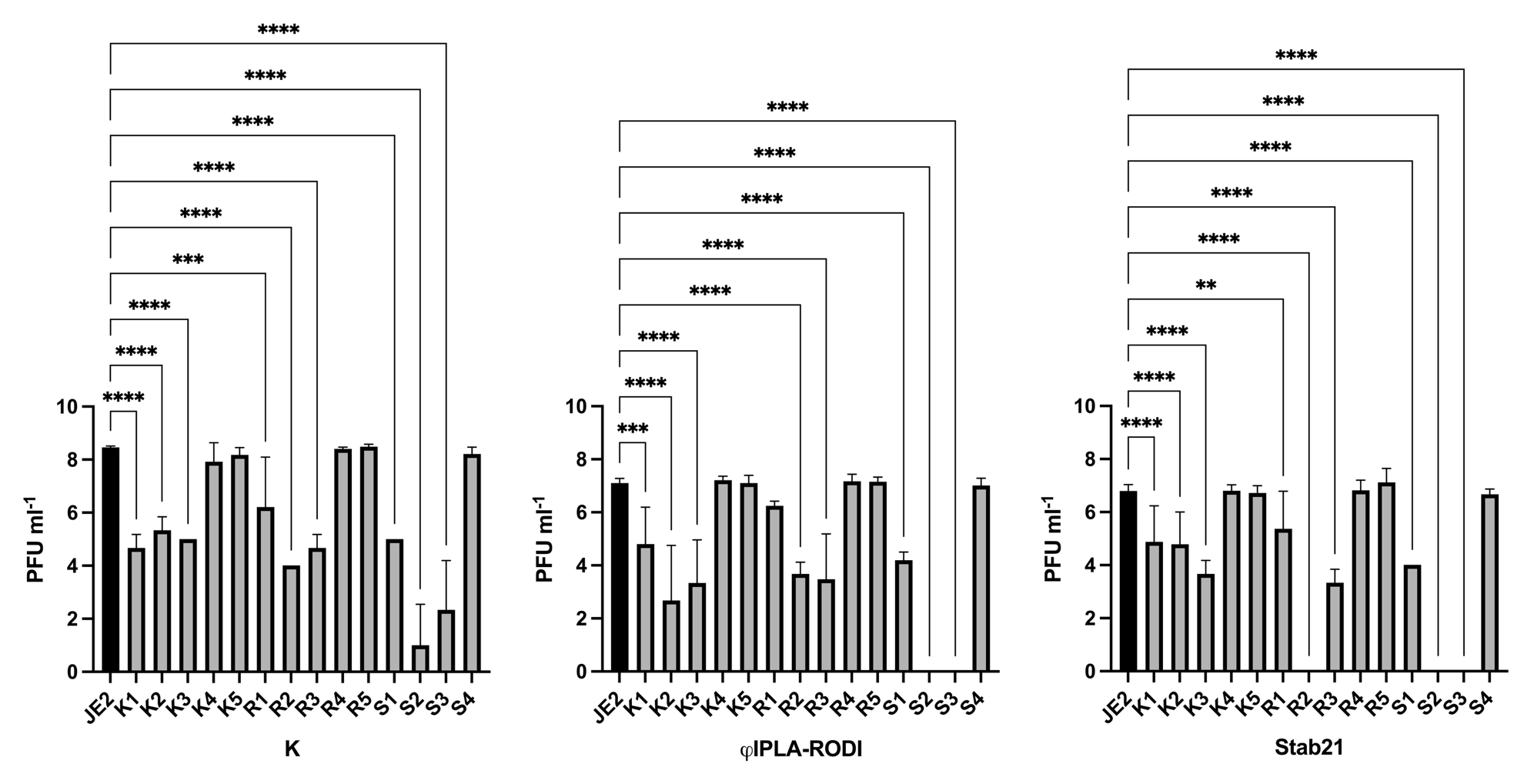

**Figure S1. Phage titer screen of potential phage resistant clones. A**. PFU ml^-1^ of phage K **B.** ϕIPLA-RODI, and **C.** Stab21 for suspected phage resistance colonies compared to the parental JE2. Data show mean with standard deviation for 6 biological replicates that was log transformed. Ordinary one-way ANOVAs with Dunnett’s multiple comparisons test were performed comparing clones to the parental JE2. Significant differences are indicated by one (*p* <0.05), two (*p* <0.01), three (*p* <0.001) or four (*p* <0.0001) asterisks (*). Clone names indicate which of the phages was used in the resistance development protocol (K for phage K, R for ϕIPLA-RODI, S for Stab21).

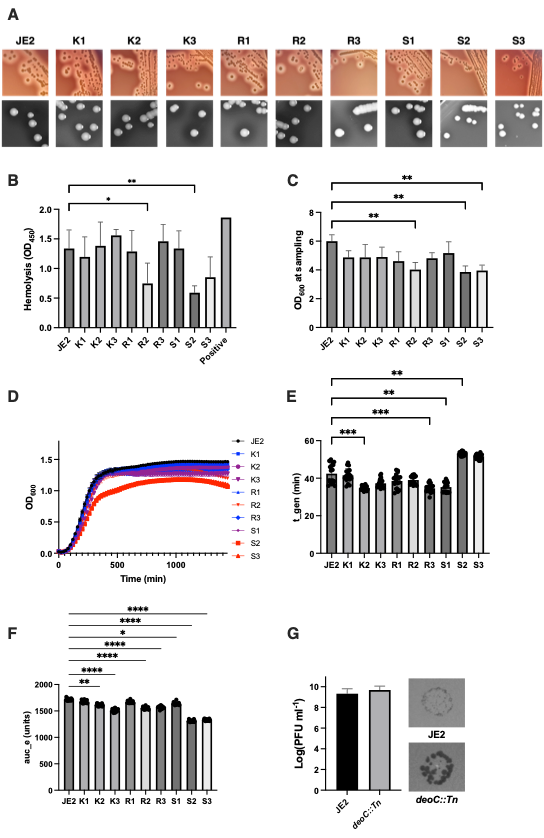

**Figure S2. Hemolysis quantification and growth phenotypes for the phage-resistant clones. A.** Colony morphology and hemolysis. **B.** Quantitative hemolysis assay for the parental JE2 and phage resistant clones. **C.** OD_600_ of the bacterial sample used for the hemolysis assay at time of sampling. **D.** Growth curves for JE2 parental strain and phage resistant clones**. E.** Generation time (t_gen). **F.** Area under the curve (empirical). For B, C, E and F, ordinary one-way ANOVAs were performed, comparing each clone to the parental JE2, with Dunnett’s multiple comparisons test. Significant differences are indicated by one (*p* <0.05), two (*p* <0.01), three (*p* <0.001) or four (*p* <0.0001) asterisks (*). Data shows 3 biological replicates, with 6 technical replicates. **G. Left**, Log transformed PFU ml^-1^ for JE2 wt and JE2 *deoC1::Tn*. Data shows 4 biological replicates. Welch’s t test showed no significant differences. **Right**, phage plaque phenotypes.

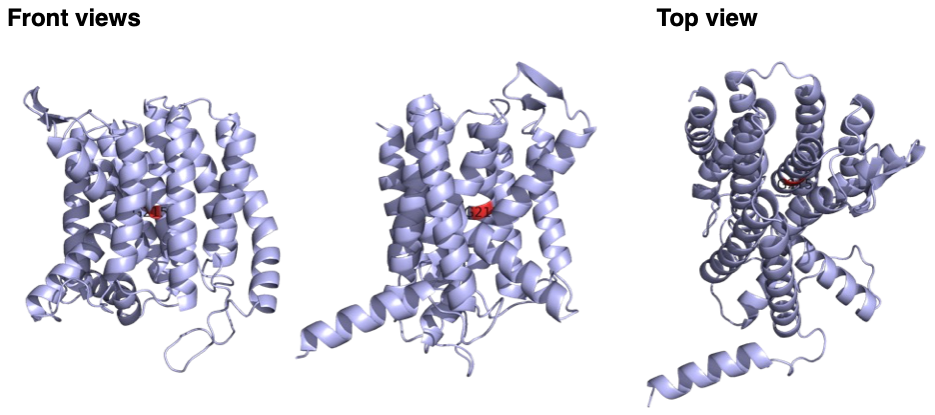

Figure S3. ‘**Front views’**, two different faces of the predicted TagO structure, ‘**Top view**’ of predicted TagO structure. G215 is labelled and indicated in red, this was mutated from glycine to arginine in clone S2 (G215R) changing the side chain from nonpolar and small, to a large, positively charged polar side chain.

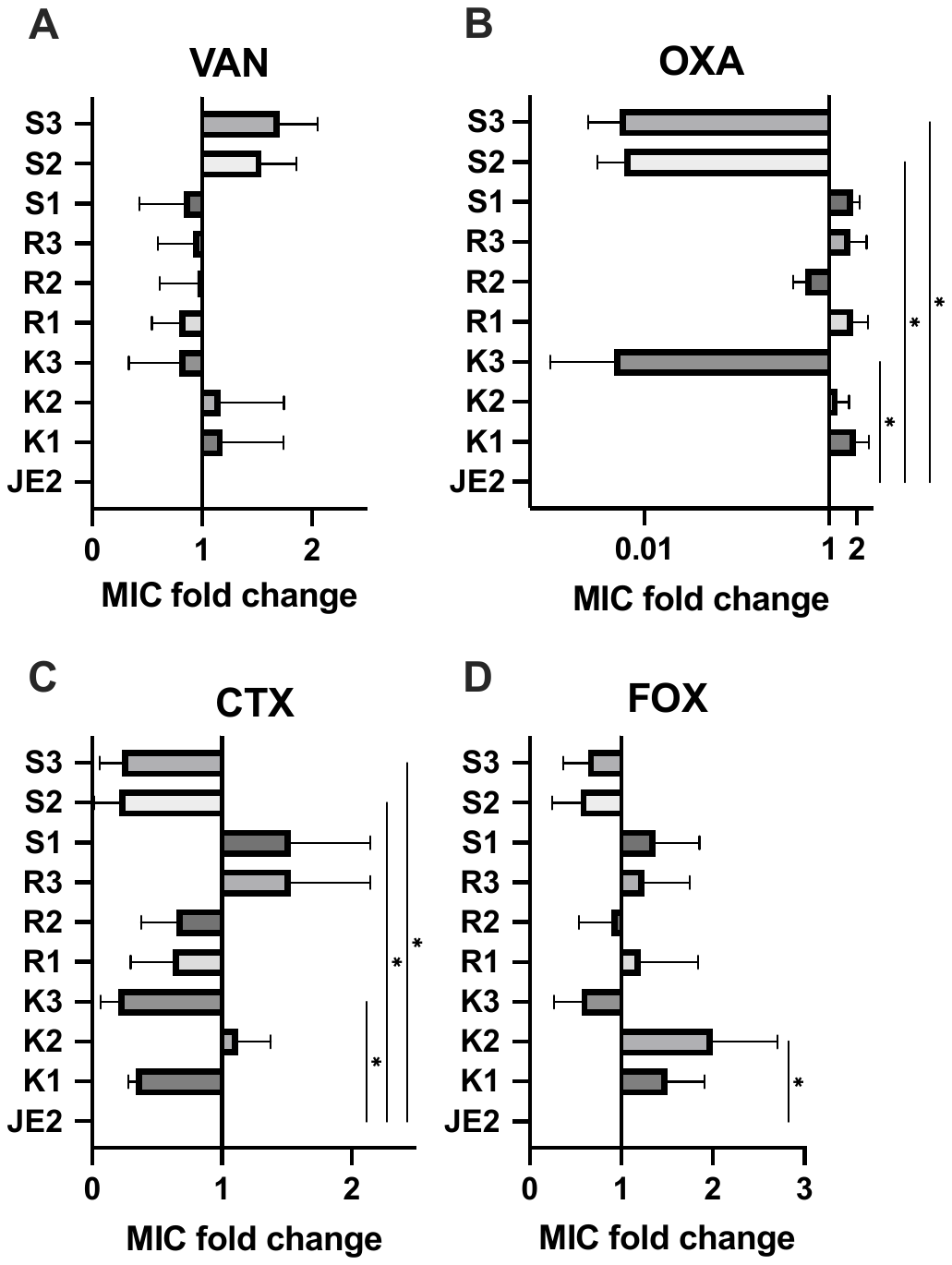

**Figure S4. Fold change of Etest MIC measurements.** Fold change calculations for Figure 3 MIC data, comparing phage resistant clones to parental JE2 for **A** vancomycin (**VAN)**, **B** oxacillin (**OXA**), **C** cefotaxime (**CTX**), and **D** cefoxitin (**FOX**). Ordinary one-way ANOVAs with Dunnett’s multiple comparisons test were performed. Significant differences are indicated by one (*p* <0.05), two (*p* <0.01), three (*p* <0.001) or four (*p* <0.0001) asterisks (*).

1.	Oduor JMO, Kiljunen S, Kadija E, Mureithi MW, Nyachieo A, Skurnik M. Genomic characterization of four novel Staphylococcus myoviruses. Archives of Virology. 2019;164(8):2171-3.

2.	Kennedy AD, Otto M, Braughton KR, Whitney AR, Chen L, Mathema B, et al. Epidemic community-associated methicillin-resistant Staphylococcus aureus: Recent clonal expansion and diversification. Proceedings of the National Academy of Sciences. 2008;105(4):1327-32.

3.	Slavetinsky J, Lehmann E, Slavetinsky C, Gritsch L, van Dalen R, Kretschmer D, et al. Wall Teichoic Acid Mediates Staphylococcus aureus Binding to Endothelial Cells via the Scavenger Receptor LOX-1. ACS Infectious Diseases. 2023;9(11):2133-40.

4.	Kreiswirth BN, Löfdahl S, Betley MJ, O'Reilly M, Schlievert PM, Bergdoll MS, et al. The toxic shock syndrome exotoxin structural gene is not detectably transmitted by a prophage. Nature. 1983;305(5936):709-12.

5.	Novick R. Properties of a Cryptic High-Frequency Transducing Phage in Staphylococcus aureus. VIROLOQY. 1967;33(1):155-66.

6.	Barbuti MD, Lambert E, Myrbraten IS, Ducret A, Stamsas GA, Wilhelm L, et al. The function of CozE proteins is linked to lipoteichoic acid biosynthesis in Staphylococcus aureus. mBio. 2024;15(6):e0115724.

7.	Monk IR, Tree JJ, Howden BP, Stinear TP, Foster TJ. Complete Bypass of Restriction Systems for Major Staphylococcus aureus Lineages. mBio. 2015;6(3):e00308-15.

8.	Myrbraten IS, Stamsas GA, Chan H, Morales Angeles D, Knutsen TM, Salehian Z, et al. SmdA is a Novel Cell Morphology Determinant in Staphylococcus aureus. mBio. 2022;13(2):e0340421.

9.	Liu X, de Bakker V, Heggenhougen MV, Marli MT, Froynes AH, Salehian Z, et al. Genome-wide CRISPRi screens for high-throughput fitness quantification and identification of determinants for dalbavancin susceptibility in Staphylococcus aureus. mSystems. 2024;9(7):e0128923.

10.	Gutiérrez D, Vandenheuvel D, Martínez B, Rodríguez A, Lavigne R, García P. Two Phages, phiIPLA-RODI and phiIPLA-C1C, Lyse Mono- and Dual-Species Staphylococcal Biofilms. Applied and Environmental Microbiology. 2015;81(10):3336-48.

11.	O'Flaherty S, Ross RP, Meaney W, Fitzgerald GF, Elbreki MF, Coffey A. Potential of the Polyvalent Anti-Staphylococcus Bacteriophage K for Control of Antibiotic-Resistant Staphylococci from Hospitals. Applied and Environmental Microbiology. 2005;71(4):1836-42.

12.	Driguez P-A, Guillo N, Rokbi B, Mistretta N, Talaga P, inventors; Sanofi Pasteur, assignee. Immunogenic Compositions Against S. aureus. WO patent WO 2017/064190 A1. 2017 2016/10/13.

**References**
